# DNN-Assisted Statistical Analysis of a Model of Local Cortical Circuits

**DOI:** 10.1101/2020.05.14.095703

**Authors:** Yaoyu Zhang, Lai-Sang Young

**Affiliations:** School of Mathematics, Institute for Advanced Study, Princeton, NJ, 08540, USA; School of Natural Sciences, Institute for Advanced Study, Princeton, NJ, 08540, USA; Courant Institute of Mathematical Sciences, New York University, New York, NY 10012, USA

## Abstract

This paper is about the use of Deep Neural Networks (DNN) to assist in the statistical analysis of a network of a few hundred integrate-and-fire neurons intended to model local circuits in the cerebral cortex. Using training data produced by direct numerical simulations, our first task was to discover, with the aid of a DNN, the mapping that yields model response for each set of parameters and input values. After evaluating the performance of the DNN surrogate both in the accuracy of its outputs and in its performance in parameter tuning, we analyzed the outputs of the well-trained DNN to gain insight into local circuits as basic cortical computational units. Because the DNN surrogate computed with vastly higher speeds than actual simulations of the neuronal network, we were able to sample large sets of parameters and input values to produce a broad statistical picture of input-output relations. One of the aims of this paper is to demonstrate that statistical analyses of this kind can provide general theoretical information on model behavior as well as suggest cortical mechanisms. Among our results are the following: Through a derivative analysis of model responses we identified a certain dichotomy in the behavior of I-neurons, leading to a characterization of high gain states which in turn offered insight into mechanisms for surround suppression. A second-derivative analysis revealed limitations of models of integrate-and-fire neurons, namely their inability to replicate the nonlinearities in gain curves observed in real neurons.

**Author summary:** Local circuits are basic computational units of the cerebral cortex, and statistical analysis of detailed models of such circuits will shed light on cortical computation. Because analytical tools are either overly simplistic or not applicable, systematic numerical exploration of high-dimensional parameter space is not feasible and simulation-based arguments are seldom more than heuristic, we propose in this paper a data-driven DNN-assisted approach that falls into the general framework of surrogate-based modeling. We consider as a test case a network of integrate-and-fire neurons intended to model local circuits in cortex. With the help of an accurate yet extremely efficient DNN surrogate, we reveal the statistics of model response, providing a detailed picture of model behavior. The information obtained is both general and of a fundamental nature, with direct application to neuroscience. Our results suggest that the methodology proposed can be scaled up to larger and more complex neuronal models.

## Introduction

Models of neuronal circuits vary widely in complexity, from firing rate models that track only two state variables, namely the averaged excitatory and inhibitory spike rates of the neurons within the circuit (see [1] for an introduction to the subject), to more sophisticated models that focus on bulk quantities as proposed by Wilson and Cowan [2], to homogeneously connected models of integrate-and-fire neurons as studied in e.g. [3, 4], to the neurofield models of [5], and to large-scale models that take into account the behaviors of individual neurons as pioneered by [6] and further developed in e.g. [7]. All of these and many other types of models – too many to enumerate – serve useful purposes but they are not without challenges: The more coarse-grained a model, the more it is amenable to analysis, but such models are not designed to incorporate neuroanatomy or capture emergent behaviors, i.e., behaviors that occur as a result of the dynamical interaction of the neurons in the circuit. At the other end of the spectrum, detailed models must necessarily involve large numbers of parameters representing unknown quantities to be determined. In existing models these parameters are either inadequately constrained or tuned by hand, a rather laborious process. As of now there is no systematic way to tune parameters in large-scale circuits, though there have been recent attempts to do that for individual neurons; see e.g. [8, 9]. The large numbers of parameters present together with the nonlinear nature of circuit dynamics has led to great difficulties in theoretical analysis on the circuit level.

On the other hand, while detailed models of neuronal networks are high-dimensional, complex dynamical systems, there is a finite number of observations that are of special importance in neuroscience. These observations include inputs to the model, transfer between regions within the model, and model outputs, mostly in the form of spike rates and currents. One may also be interested in quantitative measures that capture firing patterns (such as power spectral densities and degrees of synchrony). An important part of model analysis is to discover *input-output mapping* where, for example, input values can include model parameters and input data to the circuit and output values are one’s chosen quantities of interest. Still, studying this mapping systematically is very difficult because of the high dimensional parameter space for exploration and time-consuming simulation of each trial. To overcome the difficulties, we propose a data-driven DNN-assisted surrogate modeling approach novel in neuroscience, which is especially suited to the quantitative statistical study of the mapping behavior.

In recent years, Deep Neural Networks has achieved huge success in many areas of applications [10, 11], and its potential for fundamental research is under active exploration [12]. As an attempt in neuroscience, our basic idea is to train a deep neural network (DNN) from limited mapping data obtained by simulation as a substitute for parameter exploration via direct simulations. After learning, a well-trained DNN can serve as a surrogate model for the original neuronal circuit model to inform on output values for given sets of parameters and inputs. Because the DNN can generate input-output pairs far more quickly than actual simulations of the network model, with speeds exceeding easily 10,000 times that of actual simulations (e.g. 0.1 ms *versus* seconds to hours per trial), it has the capability to provide large collections of data points, which can then be used for systematic statistical analyses leading to a better understanding of network behavior. On the practical level, such a surrogate model can be used for automated parameter tuning in model construction, and it can be used to inform on the limitations of existing models, i.e., whether or not a model has the capability to produce certain outputs. Both model building and their statistical analysis are essential steps towards a better understanding of cortical mechanisms.

To put these ideas into a broader context, we remark that our DNN-assisted approach falls into the general framework of surrogate-based modeling, a well established practice in engineering with wide applications to many problems that involve complex simulations or experiments (see [13–15] for reviews). In biology, the use of surrogate models, e.g., support vector machines, has been recently explored in hemorrhage and renal denervation [16], and yeast mating polarization [17]. A purpose of this paper is to further promote this approach in biological modeling by demonstrating its efficacy with the assistance of a DNN in neuronal systems.

As a test case, we will in this paper apply the ideas above to a homogeneously connected integrate-and-fire neuronal network resembling a local circuit of the visual cortex. The visual cortex is an ideal place to start both because of the large amount of experimental data available and because it offers a window into many other parts of the cerebral cortex, where local circuits are believed to have similar properties. Thus this work can be seen as an attempt to gain a deeper understanding of local circuits as a basic cortical computational unit. While DNN is not the only option for the present problem – many other machine learning methods will perform as well – we anticipate applying this approach to much more complex biological networks for which the flexibility of DNN may be an advantage.

## Results

This paper is about the use of a DNN-surrogate to assist in the analysis of model outputs for a neuronal network intended to model local circuits in the cerebral cortex. The model is a network of conductance-based integrate-and-fire neurons and is described in detail in **Materials and methods** (I&F neuronal model). The deep neural net that will serve as surrogate for this model is described in **Materials and methods** (DNN surrogate). We begin by framing the problem and outlining our approach, to give the reader a sense of our perspective. This is followed by preliminary information on the capabilities of the DNN. We then present our first key results, which consist of a statistical analysis of the derivatives of model responses and their interpretation. We will demonstrate that such analyses can have surprisingly rich implications. The last part of this paper discusses another use of surrogates in biological modeling, namely to assist in the evaluation of the capabilities and limitations of models.

### DNN-assisted approach: setup and overview

We study a neuronal model of local cortical circuits with the goal of understanding its dependence on parameters and input values, and our approach is to first discover the mapping

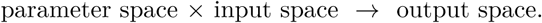

This mapping is then used to assist in the analysis of model dynamics and cortical mechanisms. The proposed methodology avoids parameter tuning, and represents a different viewpoint than standard dynamical systems approaches. As we will show, it is well suited for data-driven inferences using neural networks, and provides useful statistical information that has the potential to help unravel what goes on in complex dynamical systems.

As illustration of this methodology, we consider a homogeneously connected network of integrate-and-fire (I&F) neurons that can be thought of as a generic model of a local neuronal population. This is a dynamical system of medium complexity, with 𝒪(10^3^) state variables. The equations governing its dynamical evolution are given in

#### Materials and methods (I&F neuronal model)

The undetermined parameters of this model are the coupling weights between Excitatory (E) and Inhibitory (I) neurons. These synaptic coupling weights are denoted by *S*^*XY*^ where *X, Y* ∈ {*E, I*}; *S*^*EI*^, for example, represents the amount of influence an I-spike has on a postsynaptic E-cell. The inputs to the model network are described by the following three numbers: *η*^ext,*E*^ and *η*^ext,*I*^ are the amounts of external drive supplied to the E and I-neurons in the model population, and *η*^amb^ is an “ambient” drive intended to depict modulatory influences from outside of the population.

The objects of our study are population mean firing rates, the most fundamental statistical quantities of a neuronal circuit. Specifically, we will focus on *r*^*E*^ and *r*^*I*^, the mean firing rates of E and I-neurons in the model.

In the setup above, the mapping to be discovered and analyzed is

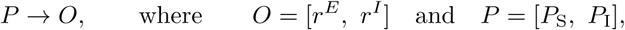

and *P*_S_ and *P*_I_ are as follows: For reasons to become clear we have chosen to represent the parameters corresponding to synaptic coupling between E and I-cells as

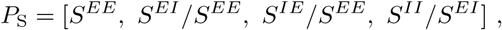

i.e., we scale the other three parameters to *S*^*EE*^ or *S*^*EI*^, and to represent the input parameters as

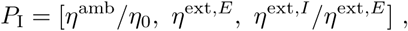

where *η*_0_ is a kind of normalizing constant.

Additionally, we specify in advance a physiological domain 𝒫 for *P*. These parameter ranges correspond to *a priori* biological constraints either deduced from indirect experimental measurements or gleaned from previous modeling results (such as [7]); they are effectively educated guesses. We also identify a physiological domain 𝒪for *O* consisting of firing rates observed in the laboratory under a variety of circumstances. We did not know in advance – and do not assume – that *P* ∈ 𝒫 will produce *O* ∈ 𝒪.

This completes a description of the setup for the rest of this paper. The mapping *P* → *O* is the mapping alluded to at the beginning of this section. We will train a DNN, details of which are given in **Materials and methods** (DNN surrogate), to learn this mapping from limited data obtained from simulation. Through the training, the DNN gradually interpolates the discrete data by a smooth function, allowing efficient evaluation and differentiation. Once we are satisfied that the DNN is performing satisfactorily, we will replace the original neuronal model by the DNN. The DNN surrogate is *a model of the original neuronal model*, one that is more limited in scope (it is focused solely on the mapping *P* → *O*) but computes at vastly higher speeds and performs efficiently certain operations that are difficult or impossible via simulation of the original model. It serves as a compass, enabling us to explore more systematically model responses as parameters are varied in a high dimensional space.

In computational modeling, DNN surrogates can assist by offering baseline values to initialize searches and by proposing parameter corrections along the way. It provides a general description of input-output relations as well as statistical information on the effects of perturbations, tasks that are well suited to the DNN. This paper is not a modeling paper and we will not get into specific instances of parameter tuning, but as an example of the theoretical insight that DNN surrogates can offer, we will present a derivative analysis of the *P* → *O* mapping. To our knowledge such an analysis has not been done before for a large network of integrate-and-fire neurons.

Finally, there are two aspects of model analysis that we would like to illustrate in this paper. One is what the model can tell us about neural mechanisms, that is, having skipped over the dynamical process, how we can now use the *P* → *O* mapping to deduce what may be going on in the neuronal model, in the hope of shedding light on what goes on in real cortex. But there is another aspect to model analysis that is also very important: all models are limited in scope because they are orders or magnitudes simpler than the real brain, and it is important to understand the limitations of a model, whether it has the capability to reproduce specific types of neural phenomena. We will finish by presenting an example of that.

### Performance of DNN surrogate

Firing rates can be measured experimentally using electrophysiology, or estimated using various kinds of optimal imaging techniques. On the theoretical level, however, how firing rates depend on network properties and inputs is not well understood, as firing rates cannot be computed analytically in semi-realistic network models such as the one described in **Materials and methods** (I&F neuronal model). In this paper we will use the DNN surrogate as an investigative tool to study these questions, but before we do that, we need to first confirm the viability of our physiological range 𝒫 (see **Materials and methods**, I&F neuronal model for details) and document the performance of the DNN surrogate. With regard to the latter, we will examine the accuracy of the DNN surrogate as a function of the size of its training set, and we will investigate its performance in parameter tuning, i.e. to solve the inverse problem of locating parameters to produce target outputs.

#### Viability of parameters and DNN performance

To confirm the viability of our *a priori* choice of physiological domain 𝒫, we randomly selected 20000 sets of *P* from this domain and computed from simulations their mean population firing rates *O*, which forms a training dataset 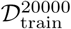. The results are presented in Fig. 1(a). The physiological domain 𝒪 consists of values in the region bounded by the trapezoid. Fig. 1(a) confirms that parameters from 𝒫 produce firing rates in a broad region containing 𝒪, justifying our choice of 𝒫. It also shows that only about 10% of the outputs *O* actually fall into the trapezoidal zone, underscoring the challenges in prescribing *P* for desired firing rates.

**Fig 1.**
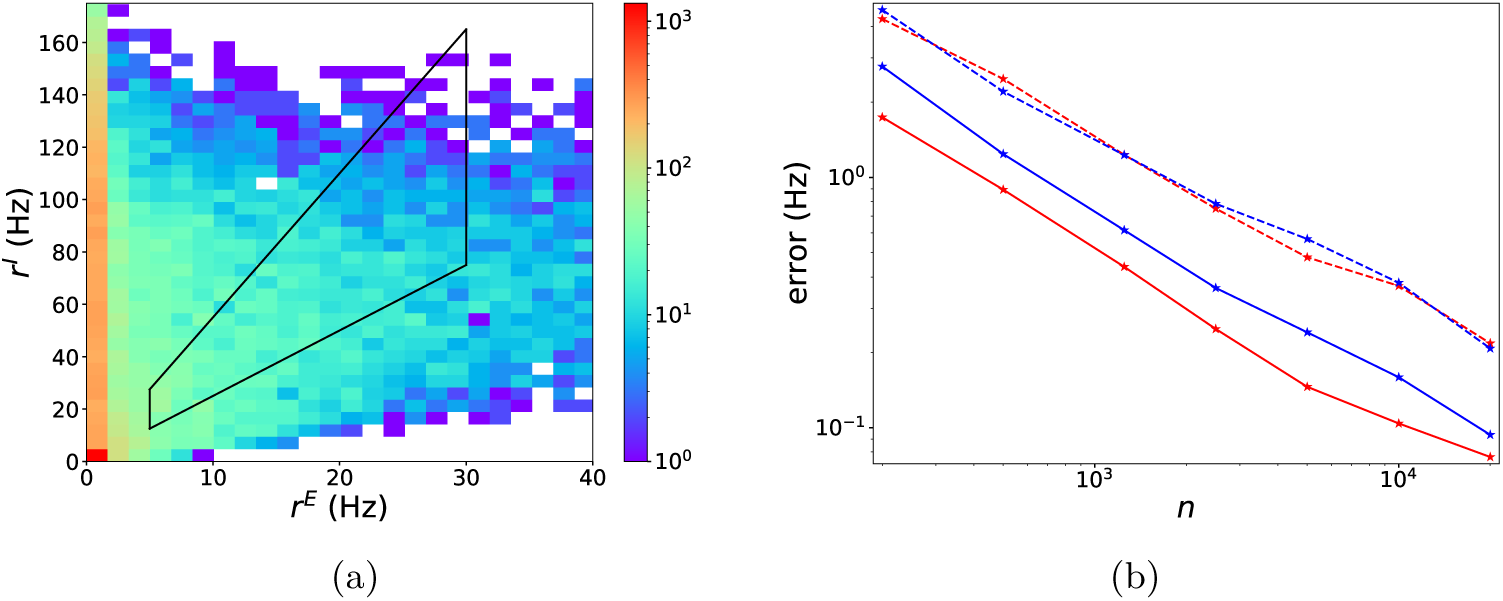
Viability of parameters and DNN performance. (a) Heat map of frequency of *O* = [*r*^*E*^, *r*^*I*^] in 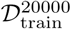. (b) Testing accuracy of DNN well-trained on 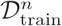 as a function of *n*. Mean-absolute errors (MAE) (dasheded line) and root-mean-square errors (RMSE) (solid line) for *r*^*E*^ (red) and *r*^*I*^ (blue) are exhibited.

We then investigated the accuracy of DNN surrogates trained on datasets of various sizes from 200 to 20000, The mean-absolute error (MAE) and root-mean-square error (RMSE) of well-trained DNNs on the testing dataset are presented in Fig. 1(b). The error follows approximately a power law decay of ∼*n*^−2*/*3^ where *n* is the size of the training set, much faster than the ∼*n*^−1*/*7^ law implied by the curse of dimensionality. This curse-of-dimensionality free convergence behavior of DNN is supported by theoretical studies [18, 19]; it is one of the reasons why DNNs are widely used for high dimensional problems.

Note also that with a surprisingly small size of 500 training data points, a small error (MAE) of ∼1Hz was obtained. In these experiments, errors are roughly independent of firing rate, resulting in smaller relative errors at high firing rates and larger relative errors at low firing rates. For predictions that result in a target E-firing rate of ∼10Hz, the relative prediction errors of our DNNs typically are ∼10% and ∼1% with 500 and 20000 training data, respectively. By theoretical studies of DNN [20, 21], such a good performance suggests a low complexity/frequency nature of the *P* → *O* mapping.

The sigmoid function 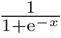, which is used as the activation function of our DNN, yields far lower testing errors than the popular choice of ReLU. A key difference between ReLU and sigmoid activation is their smoothness, a property more important for regression problems as considered in this paper than for classification problems which are commonly considered by the AI community. As suggested in Ref. [21], when the smoothness of activation matches the smoothness of the target function, an optimal error bound can be achieved. Thus the better empirical performance of sigmoid compared to ReLU activation suggests a smooth nature of the *P* → *O* mapping, a point we will revisit later on in our analysis. We remark also that smooth activation functions like sigmoid or tanh (hyperbolic tangent, a rescaled sigmoid function) have been shown to be better choices for other regression problems, e.g., in molecular dynamics simulation [22, 23].

In the rest of this paper, we will use the most accurate DNN well-trained on 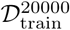 as a surrogate to investigate the statistical properties of the *P* → *O* mapping.

#### Performance of DNN surrogate for parameter tuning

Realistic models of neuronal circuits typically involve large numbers of parameters corresponding to quantities not directly measurable in the laboratory. Fitting these parameters to experimental observations is an essential task. Up until now, this task has often been done “by hand”, relying on the experience of the modeler. As such, it is both laborious and time-consuming if it can be successfully carried out at all. Because of the high dimensionality of the parameter space, and the difficulty in directly computing the derivatives ∇_*P*_ *O* from discrete data points, automated gradient-based approaches widely used in many applications are not viable in this kind of parameter tuning.

Our first demonstration of the usefulness of an accurate DNN surrogate is to apply it to the problem of automated parameter tuning. This is an inverse problem, requiring that we find parameter *P* given target output *O*_target_. Assisted by the DNN surrogate *Ô*(*P*) well trained on 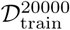, whose derivatives can be easily computed by back-propagation, a gradient-based approach can be efficiently applied as follows. In each iteration step *t, P*^*t*^ is updated as

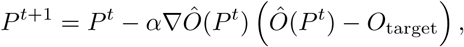

where *α* is the learning rate.

Figure 2 shows the results of a numerical experiment we performed. In this experiment, the initial parameter *P* ^0^ was randomly sampled from 𝒫, and if *P*^*t*^ fell outside of the domain, it was projected back to 𝒫. For each *O*_target_, we selected 100 random initial parameters. After 10000 steps of iteration, all final *P* ‘s with predicted output sufficiently close to the target, e.g., with ‖*Ô*(*P*) − *O*_target_‖_1_ *<* 0.2Hz, formed the candidate set of parameters for *O*_target_. For acceleration, we incorporated the scheme of Adam [24].

**Fig 2.**
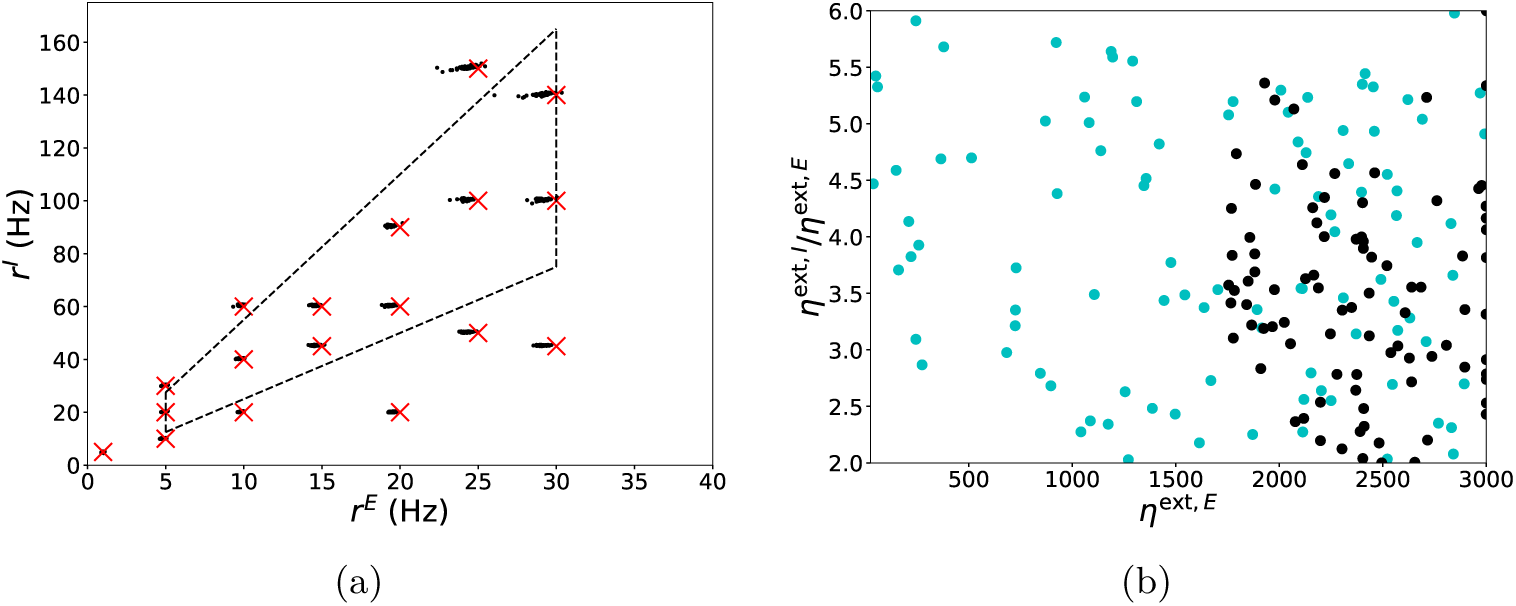
Visualization of DNN-assisted parameter tuning. (a) Visualization of accuracy of parameter tuning. Crosses are target firing rates, black dots are the simulated output of the candidate parameters found through gradient-based approach assisted by the DNN surrogate. The trapezoid constrained by the dashed lines indicates the physiological domain of output. (b) Parameters at initialization (cyan dots) and after tuning (black dots) for target (25, 100)Hz projected to the 2D plane of *η*^ext,*E*^ and *η*^ext,*I*^*/η*^ext,*E*^.

In general, given *O*_target_, a randomly chosen parameter in 𝒫 has probability *<* 0.01% to be a candidate parameter. The iterative scheme above was intended to autonomously steer it towards a candidate.An example is depicted in Fig. 2(b): parameters at initialization (projected from 7D) are represented by cyan dots; they are steered to black dots through the tuning for a given target. For each target, we found that of the 100 initial parameters picked, on average over 90% successfully yielded a candidate after 10000 steps. The accuracy of the candidate parameters were then evaluated by comparing their simulated outputs (black points) with the corresponding targets (crosses). The error was larger than the mean testing error of ∼0.1Hz for the DNN, as can be expected for an inverse problem. However, except from parameters in the periphery of 𝒫, most tuning results were faithful to the target. Note that, the accuracy of the above parameter tuning approach can be further improved by incorporating a few trials of simulation online to fix the local prediction error of the DNN surrogate.

Fig. 2(b) illustrates another important point, namely that for a given target *O*_target_, the parameters obtained by the above tuning process are far from unique. In Fig. 2(b), different pairs of input strengths *η*^ext,*E*^ and *η*^ext,*I*^ indicated by black dots (each with its own accompanying parameters in the other 5 dimensions) give rise to the same (E,I)-firing rates of (25, 100)Hz*}*. Indeed, if the *P* → *O* mapping is smooth, one would expect, for each given *O*_target_, the set {*P* : *Ô*(*P*) = *O*_target_} to be a 5D submanifold in our 7D parameter space. In modeling, additional physiological phenomena will likely place further constraints on the set of viable parameters.

### Statistical analysis of parameter dependence: first derivatives

Crucial for understanding cortical mechanisms is a quantitative description of how the firing rates of a brain region depend on its structural and input parameters. Yet except for extremely idealized models with few state variables, there is no explicit relation between these parameters and firing rates, and exploration of parameter space via simulations is not feasible as we have explained earlier. In this paper, we propose a statistical approach to this problem via the use of DNN surrogates.

In Fig. 1(a), we presented the statistics of firing rate responses for parameters in 𝒫. This section focuses on statistics on the *derivatives* of output responses. Our study is assisted by the well-trained DNN surrogate *Ô*(*P*), which allows very efficient evaluation and differentiation. To our knowledge, this is the first time that parameter dependence of firing rates in integrate-and-fire models are systematically investigated through a statistical analysis.

Quantitative information on ∇_*P*_ *O* will shed light on a number of questions. Of particular interest is a system’s response to changes in its input. As we will show, our statistical analysis points to a dichotomy in the response behavior of neuronal populations. It supports a novel interpretation of “high gain” that may have implications in cortical phenomena such as surround suppression.

#### Derivative analysis

Recall that in our model, input parameters are

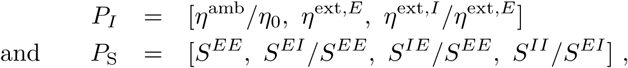

and output parameters are

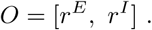

Using the DNN surrogate *Ô*(*P*) trained on 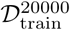 (see **Materials and methods**, DNN surrogate), one can easily compute ∇_*P*_ *Ô*, which approximates ∇_*P*_ *O*, over a very large number of input parameters. Fig. 3(a) and b show the distributions of partial derivatives 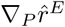 and 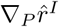, respectively, with respect to each of the seven parameters in {*P*_*S*_, *P*_*I*_}. (We write 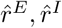 to stress that these results are computed from the DNN surrogate *Ô*(*P*).) The histograms in Fig. 3 were computed from 5 *×* 10^5^ randomly selected *P* ∈ 𝒫, keeping only the ∼10% of *P* for which *Ô*(*P*) ∈ 𝒪 and discarding the rest.

**Fig 3.**
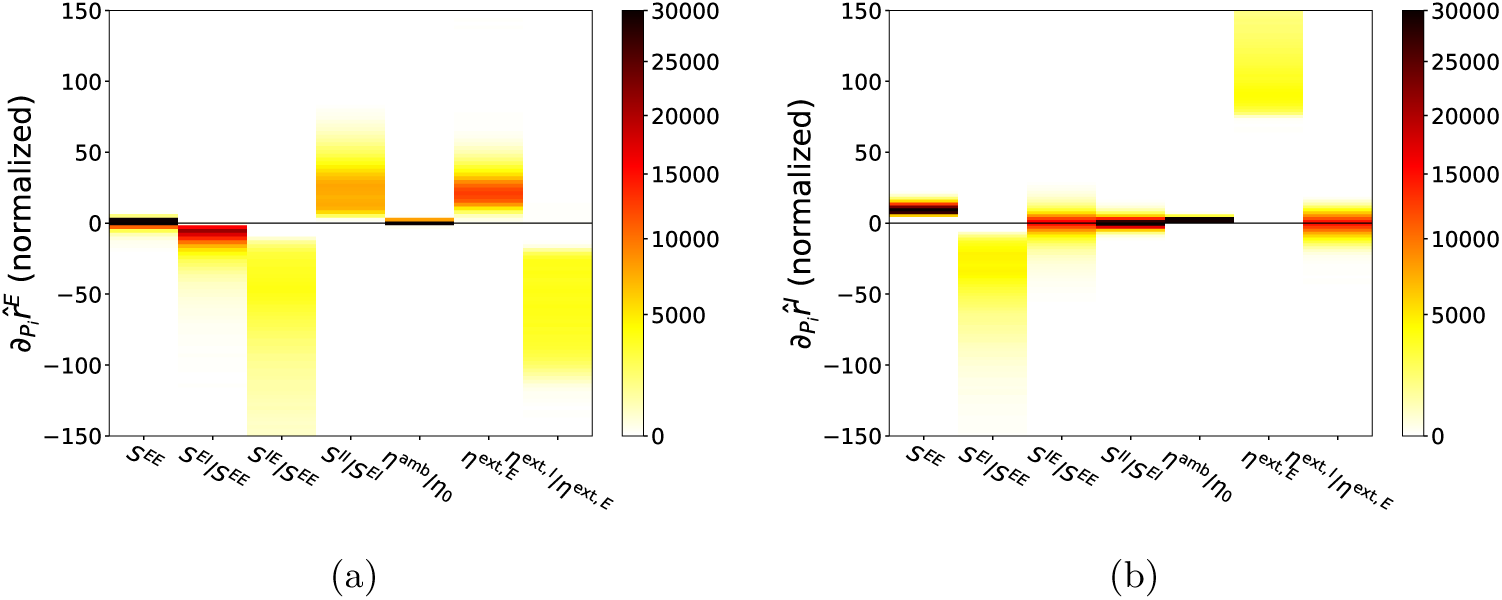
First derivative statistics. 2D Histograms of ∇_*P*_ *Ô* for *P* ‘s satisfying *P* ∈ *P* and *Ô*(*P*) ∈ 𝒪. 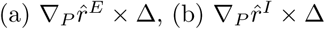 where Δ is a vector consisting of the length of physiological domain for each parameter in *P*. Δ is used to fix 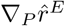 to a dimensionless unit with 𝒫 scaled as a unit box.

To familiarize the reader with the meaning of the plots in Fig. 3, consider, for example, differentiating with respect to *S*^*EI*^*/S*^*EE*^ keeping the other 6 parameters fixed. The second columns of the two panels show that both 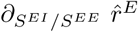 and 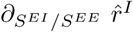 are almost always negative indicating that increase of strength from I to E consistently decreases the firing rate of both the E and the I-population. In addition, the magnitude of 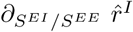 is in general larger than 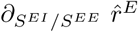, indicating that changes in *S*^*EI*^*/S*^*EE*^ have a larger effect on I-firing rate, not surprisingly since I-firing rates are generally 3 − 4 times larger than E-firing rates.

Differentiating with respect to *S*^*IE*^*/S*^*EE*^, on the other hand, yields rather curious results: while 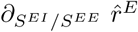 is always strongly negative, 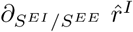 can be positive or negative with a relatively small magnitude. This statistical result suggests the existence of an interesting regime where increasing the synaptic strength of E to I (while keeping that from E to E fixed) decreases the firing of the I-population (even though the strength of E to I is increased!) and it suppresses the firing of the E-population (even though I-firing is lowered!). This model behavior is reminiscent of the “paradoxical effect” identified earlier in [25–28]. We will revisit this point in the next subsection.

The following information on the dependence of response properties on parameters can be gleaned from Fig. 3:

1. *Parameter dependences are nonlinear*. Fig. 3(a) and (b) ruled out the possibility that *r*^*E*^ and *r*^*I*^ are as simple as a linear function of *P* because most of the partial derivatives are clearly nonconstant; some in fact have quite a large spread.
2. *Dependence on η*^amb^*/η*_0_ *is insignificant and dependence on S*^*EE*^ *is weak*. As the other three synaptic weights are indexed to *S*^*EE*^ in our bookkeeping, the relatively weak dependence on *S*^*EE*^ when the other parameters are fixed confirms our conjecture (see **Materials and methods**, I&F neuronal model) that not a great deal changes when the four synaptic weights *S*^*EE*^, *S*^*EI*^, *S*^*IE*^ and *S*^*II*^ are scaled up and down together as long as they maintain the same relationship.
3. *Near-monotonicity of the function* 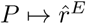. Differentiating 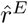, one sees that 5 out of the 7 partial derivatives have a single sign, i.e., they are either positive or negative for all the parameters tested, and the remaining two are relatively small. All this points to a simple structure for the mapping 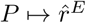. One notes also that the signs of the 5 all go in directions expected: increasing I to E and E to I lowers 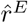 as one would expect as that increases the power of the inhibition, increasing I to I increases 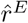, and increasing external drive to E increases 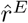 while increasing external drive to I lowers 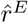 – all are as expected.
4. *The mapping* 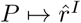 *is more complex*. Our statistics show that the I-responses are not as clean as E-responses, in that changes in 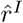 in response to increases in *S*^*IE*^*/S*^*EE*^, *S*^*II*^*/S*^*EI*^ and *η*^ext,*I*^*/η*^ext,*E*^ can be positive or negative. As noted earlier, the idea that increasing drive to I-neurons could decrease 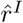 is somewhat counter-intuitive. With the help of the DNN surrogate, we examine next in more detail the circumstances surrounding this response reversal of I-neurons.

### Cortical mechanisms via DNN-assisted derivative analysis: an illustrative example

The phenomenon that stimulation of an inhibitory population not only decreases the activity of the excitatory population but that it can also decrease the activity of the stimulated population is known to the neuroscience community. The intuition is that the excitatory population is sufficiently suppressed that the total excitation received by the inhibitory population is reduced [25–30]. In rate models, it has been demonstrated mathematically that this occurs in inhibition stabilized networks (ISN), where recurrent excitation is strong and the regime is stabilized by inhibition [25–27]. Models with multiple inhibitory populations have also been investigated recently [28, 31–33]. For network models of integrate-and-fire neurons such as the one studied here, analytical approaches are not viable, and conditions for the reversal of I-response have not been investigated. This is what we would like to do using a DNN-assisted statistical analysis.

#### Response of I-neurons: a dichotomy

Following up on the observation in Item (4) above, namely that 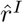 may increase or decrease in response to changes in *S*^*IE*^*/S*^*EE*^, *S*^*II*^*/S*^*EI*^ and *η*^ext,*I*^*/η*^ext,*E*^, we looked into potential correlations between the signs of these partial derivatives. The results are shown in Fig. 4(a), and they show that the signs of these partial derivatives are highly correlated to one another, with correlations very close to ±1 (see **Materials and** methods, Correlation analysis and logistic regression for details). This suggests the existence of two distinct regimes: one in which an increase in *S*^*IE*^*/S*^*EE*^ or *η*^ext,*I*^*/η*^ext,*E*^, or a decrease in *S*^*II*^*/S*^*EI*^ causes *r*^*I*^ to increase, and another in which the same changes cause *r*^*I*^ to decrease.

**Fig 4.**
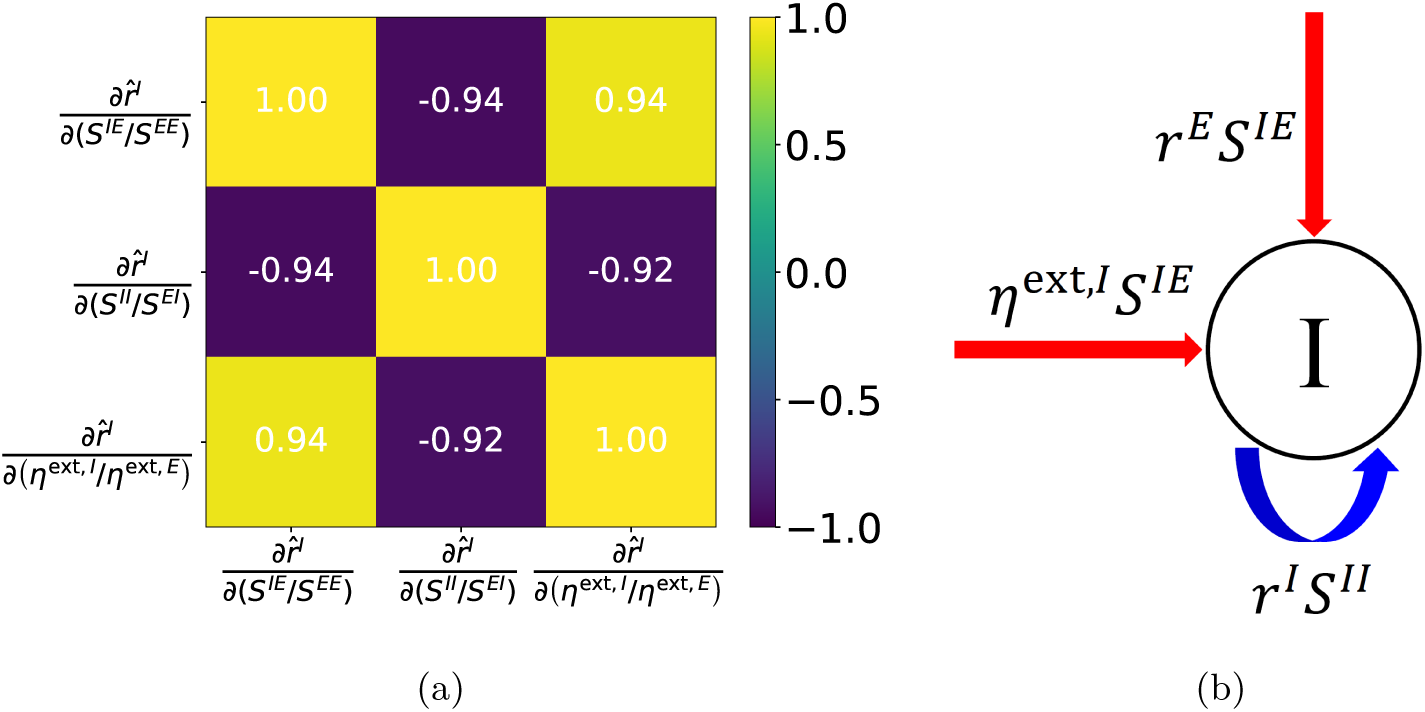
Illustration and analysis of I-population response. (a) Correlation matrix of the sign of 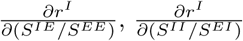 and 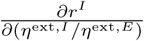. (b) Illustration for different input sources received by the I population. Red indicates excitation and blue indicates inhibition.

While *S*^*IE*^, *η*^ext,*I*^, and *S*^*II*^ directly contribute to the input received by the I-population as illustrated in Fig. 4(b), the positivity of correlations with respect to changes in *S*^*IE*^ and *η*^ext,*I*^ is not clear *a priori*, because these changes also affect the firing rates of E-neurons, and the synaptic excitatory input from within the population to an I-neuron is determined not just by *S*^*IE*^ but also by *r*^*E*^, the firing rate of the E-population. The same is true for the effect of *S*^*II*^ : increasing that does not necessarily mean that an I-neuron will receive greater suppression, because the amount of inhibitory synaptic input it receives depends also on *r*^*I*^.

To summarize, our results as shown in Figs. 3 and 4(a) show that the set of parameters

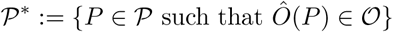

can be divided into two distinct groups according to the sign of 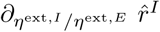, equivalently the sign of either one of the other two partial derivatives. This means that the mapping 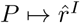, which we had noted earlier might be considerably more complex than 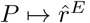, has a fairly simple structure after all. The simplicity of the mapping *P ↦O* may be the reason why DNNs achieve very good accuracy even for training datasets of small sizes.

Below we will refer to the phenomenon of 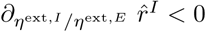 as “inhibitory response reversal”.

#### Correlating network properties to inhibitory response reversal

We first used the seven quantities in *P* to predict the sign of 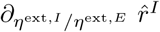 by logistic regression, i.e., we used the logistic function 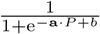 to fit the probability of sign 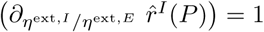 (it is either equal to 0 or to 1 in this problem). Similar to linear regression for real-valued output, in machine learning, logistic regression is often a first try for fitting binary output with real-valued input. After regression, the accuracy of prediction using the sign of 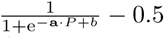 is ∼83% over *P* ∈ 𝒫^*\**^. This accuracy indicates that signs of the target can roughly be separated by a hyperplane in the space of *P* (100% indicates perfectly linearly separable while chance rate 50% indicates complex behavior far from linearly separable). Relative importance of each parameter *P*_*j*_ is evaluated by 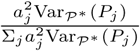, where Var_𝒫*_ indicates the variance over 𝒫^*\**^ (see Fig.5(a)).

Clearly, *S*^*IE*^*/S*^*EE*^ and *η*^ext,*I*^*/η*^ext,*E*^ are the two most salient factors for regime determination. The performance of regime separation using these two parameters is shown in Fig. 5(c). One can see a trend that smaller values of *S*^*EI*^*/S*^*EE*^ and *η*^ext,*I*^*/η*^ext,*E*^ indicating weak drives to the I-population are more likely to result in inhibitory response reversal. The prediction accuracy by logistic regression using only these two parameters yields a significantly worse accuracy of ∼67%, indicating that the ignored input dimensions in fact play nonnegligible roles in the prediction, and there is no clean linear separation between the two regimes in the space of *P*.

**Fig 5.**
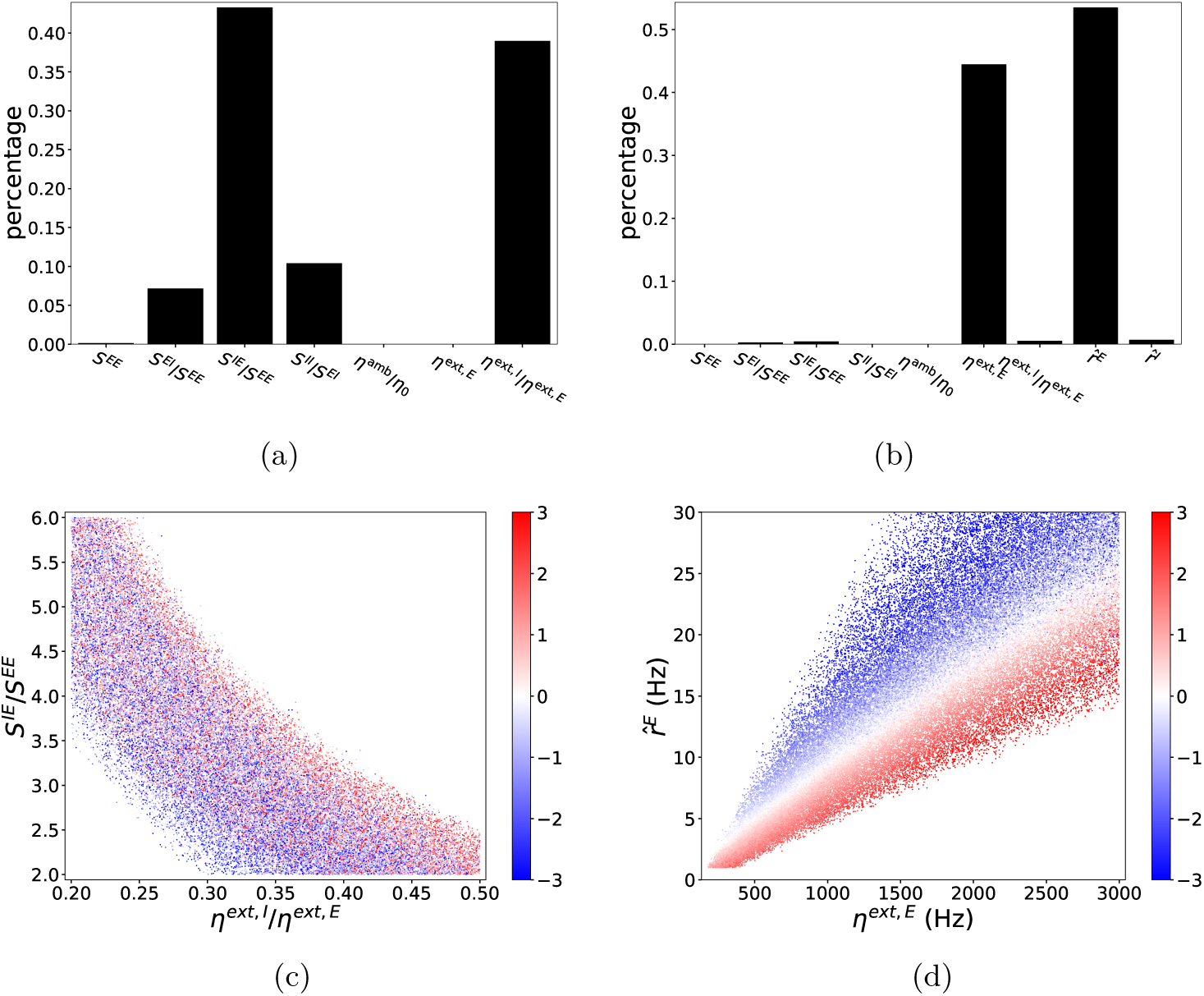
Analysis of inhibitory response reversal. Upper panels: Relative importance of each parameter in the best logistic predictor of the sign of 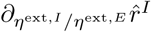 using (a) *P* only, (b) both *P* and *Ô*. Bottom panels: 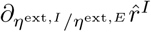 (red indicates strongly positive and blue indicates strongly negative) over different pairs of (c) *S*^*IE*^*/S*^*EE*^ and *η*^ext,*I*^*/η*^ext,*E*^, (d) 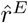 and *η*^ext,*E*^, both selected based on the importance analysis in (a) and (b) respectively.

As noted in Fig. 4(b), *r*^*E*^ and *r*^*I*^ also play important roles in determining the inputs that go into I-neurons, so we experimented next with using *P* and *Ô* together for the prediction of the sign of 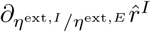. After logistic regression, we achieved a surprisingly high accuracy of ∼97%. Moreover, as shown in Fig. 5(b), *η*^ext,*E*^ and 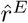 stood out as effectively the only key factors that mattered for the prediction. By using only *η*^ext,*E*^ and 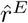, one can still achieve a very high prediction accuracy of ∼94%. This surprisingly good performance is illustrated in Fig. 5(d), where the two regimes characterized by the sign of 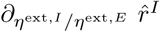 are very well separated by a line of the form 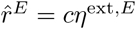 for some *c >* 0. Note that, this *c* is clearly independent of 7 model parameters, however, may depend on other factors like connectivity rates fixed in our model.

A regime with a large excitatory response to external drives can be thought of as having high gain. Our results suggest that a natural definition of *high gain* might be *r*^*E*^ *> cη*^ext,*E*^ for the critical value of *c* defined above. With this notion of gain, the above statistical analysis suggests that

##### inhibitory response reversal occurs in a regime of high gain

It is difficult to compare directly the parameters used in rate models and in networks of integrate-and-fire neurons. In our model, the physiological ranges of the parameters are chosen to be consistent with experimental data [7]. For parameters in this range, we found that sufficiently high gain, i.e., *r*^*E*^*/η*^ext,*E*^ *> c*, is the best condition for inhibitory response reversal. This finding is new, it is quantitative, and it was discovered entirely through our DNN-assisted analysis. The implications of this finding and its relation to ISN need to be explored; that will be done elsewhere. We finish with a direct application of this idea.

#### Plausible explanation for surround suppression

Surround suppression is a well documented visual phenomenon. It refers to the fact that a neuron’s sensitivity to a stimulus is modulated by the extent of the stimulus outside of its classical receptive field. The discussion below is far from a systematic study of surround suppression, which is a wide-ranging and important topic in its own right. We wish to point out only a plausible explanation for the suppression associated with spatially extended stimuli that follows from the observations above.

To briefly review the phenomenon, consider an excitatory neuron in the primary visual cortex, V1. Drifting gratings of various sizes aligned with the neuron’s orientation preference and centered at its receptive field are presented. It has been observed that while the neuron spikes vigorously in response to smaller gratings, its response peaks at a certain grating radius and decreases as the size of the grating continues to increase, leveling off eventually when the stimulus is many times the size of its classical receptive field [34]. This decrease in firing rate of a neuron at the center when the surround is also stimulated is called surround suppression. Experimental measurements of a quantity called suppression index indicates that the suppression of E-neurons can be quite strong depending on layer within V1 [35]. For some layers, firing rates for large gratings may be no more than half those for smaller gratings. A similar phenomenon has been found to hold for I-neurons, though the decline in firing rate is smaller [26].

Here is how our results may be relevant:

Consider a local population located at the center, receiving external input from feedforward and feedback sources as well as from within its own layer via long-range connections. We hypothesize that for E-neurons in this population, as the size of the stimulus increases, *η*^ext,*E*^ first increases and then saturates as the size of the grating continues to increase, whereas input to the I-population, *η*^ext,*I*^, increases for a while longer saturating at a larger grating radius. This means that *η*^ext,*I*^*/η*^ext,*E*^ is at first constant and later increases. We further hypothesize that the circuit is always in a high gain state, i.e., *r*^*E*^*/η*^ext,*E*^ is always larger than the critical value *c* defined above.

When *η*^ext,*E*^ and *η*^ext,*I*^ are both increasing and *η*^ext,*I*^*/η*^ext,*E*^ is constant, our derivative analysis asserts that both *r*^*E*^ and *r*^*I*^ should be increasing, consistent with experimental observations before the size-tuning curves peak. When *η*^ext,*E*^ saturates and *η*^ext,*I*^ continues to increase, we are in the situation where the partial derivatives with respect to *η*^ext,*I*^*/η*^ext,*E*^ becomes relevant, and if the population is in a high gain state, then our derivative analysis predicts that *r*^*I*^ would decrease though not as steeply as *r*^*E*^, a prediction in agreement with experimental data.

To summarize, our proposed explanation suggests that it is entirely possible to have both *r*^*E*^ and *r*^*I*^ decrease while *η*^ext,*E*^ and *η*^ext,*I*^ are both increasing, provided the relative rates of increase in *η*^ext,*E*^ and *η*^ext,*I*^ are as above. We have also identified the property that is the key to what makes this possible, namely that the population should be in a state of high gain.

### Analysis of second derivatives

Second derivatives reflect the acceleration and deceleration of the output in response to changes in parameters. In this section, we study the statistics of second derivatives, and investigate the model’s capability to produce nonlinear outputs in response to increasing drive.

#### Distribution of second derivatives

Fig. 6 displays the distribution of second partial derivatives of 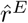 and 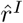 with respect to each dimension of *P*. As an example of what these histograms tell us, consider the fact that 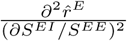 and 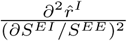 are always positive. Combined with our earlier result that 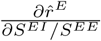 and 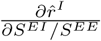 are both negative, we get the following picture: As *S*^*EI*^ increases (with all other parameters fixed), 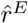 and 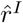 both decrease, and the graphs are convex. The effect of *S*^*IE*^ is curious: As *S*^*IE*^ increases, 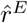 decreases and the graph is (quite strongly) convex. The graph of 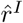 is also convex, but since 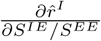 can change sign, there is the possibility that it can decrease first and later increase.

**Fig 6.**
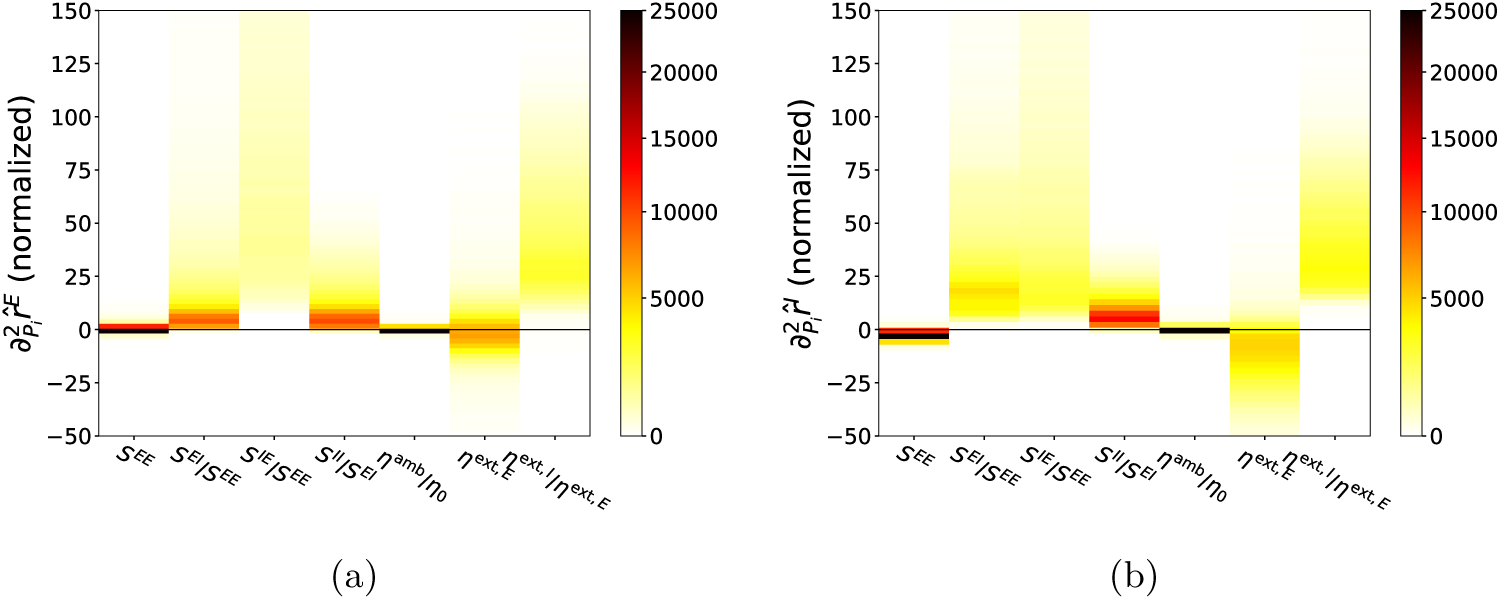
Second derivative statistics. Histograms of second order partial derivatives for the physiological parameters yielding physiological output, i.e., as *P* vary over 𝒫^*\**^ is presented. 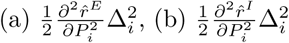 for index *i* = 1, *…*, 7, where Δ is the length of physiological domain for each parameter in *P*. Here 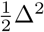 is used to fix second partial derivatives to a dimensionless unit with physiological domain of *P* scaled as a unit box.

In general, the following response properties can be inferred from the statistics of second derivatives.

1. *Outputs are not describable by second-order polynomials*. Fig. 6(a) and (b) rule out the possibility that *r*^*E*^ and *r*^*I*^ can be as simple as second-order polynomials of *P*. Most second partial derivatives are clearly nonconstant, and some have quite a wide spread.
2. *Insignificance of dependence on S*^*EE*^ *and η*^amb^*/η*_0_. This is consistent with results from our first derivative analysis.
3. *Convexity of r*^*E*^ *and r*^*I*^ *as functions of all parameters in P except for η*^ext,*E*^. This property further supports the simplicity of the mapping *P ↦ O*.
4. *Nonlinearity of gain curves*. We are concerned here with the second derivatives of *r*^*E*^ and *r*^*I*^ with respect to *η*^ext,*E*^, i.e. when both *η*^ext,*E*^ and *η*^ext,*I*^ are increasing with the ratio of *η*^ext,*I*^*/η*^ext,*E*^ fixed. Firing rates almost always increase monotonically by our first derivative analysis, but they can accelerate or decelerate as our second derivative analysis shows. A more quantitative analysis reveals the following, however: While a typical change of 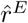 is *>* 30Hz over the input domain, the normalized second derivative 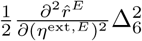 is typically between *±*10Hz. The smallness of the second derivative compared to the first suggests that gain curves are statistically more likely to be fairly linear for our model with physiological parameters.

As mentioned in the Overview of Results, one of the uses of a surrogate model is to inform on the limitations of the original neuronal network model. In real cortex, gain curves have been observed to be sigmoidal in shape. Item (4) in the second derivative analysis above raises the question of whether neurons in the model described in **Materials and methods** (I&F neuronal model) are capable of producing such nonlinear gain curves. We now investigate this question more systematically using the DNN surrogate.

#### Generation of nonlinear gain curves

Gain curves capture changes of *r*^*E*^ in response to changes in external input. For convenience, we let *P* ^−^ denote all the parameters of *P* except for *η*^ext,*E*^, and study the gain curve 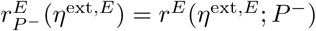. In physiological experiments, sigmoidal gain curves are often observed [36], and neurotheories hinging on the shapes of gain curves have been proposed [37]. In this section, we study with the help of the DNN surrogate whether the model described in **Materials and methods (I&F neuronal model)** is capable of producing gain curves that are sigmoidal in shape.

To capture the sigmoidal property, we require, for definiteness, that *r*^*E*^ as a function of *η*^ext,*E*^ be accelerating for *r*^*E*^ ∈ [5Hz, 15Hz], and decelerating for *r*^*E*^ ∈ [20Hz, 30Hz]. For each *P* ^−^ in the physiological range, we increase *η*^ext,*E*^, and as 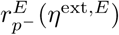 increases, we identify the intervals *J*_1_, *J*_2_ of *η*^ext,*E*^ that correspond to 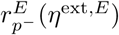 falling in [5Hz, 15Hz] and [20Hz, 30Hz] respectively. We then compute the mean values of 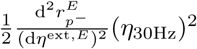 on *J*_1_ and *J*_2_, and call them *m*_1_(*P* ^−^) and *m*_2_(*P* ^−^). Here, *η*_30Hz_, which is determined by solving 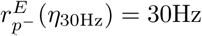, is used to normalize the second derivative to a unified dimensionless unit.

In Fig. 7, the *x* and *y*-axes show the *m*_1_ and *m*_2_ values for each plausible *P* ^−^ satisfying (i) [*P* ^−^, *η*^ext,*E*^] ∈ 𝒫 and (ii) *ÔP* − (*η*^ext,*E*^) ∈ 𝒪 for 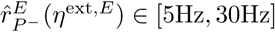. The lower right box bounded by the two black lines describes the region with the desired sigmoidal properties. As one can see, very few data points lie in this box. Some examples of gain curves are displayed in Fig. 7(b), (c) and (d), where results from the DNN surrogate and firing rates simulated directly from the neuronal network are superimposed. At least in these examples, our DNN surrogate quite accurately emulates the true behavior of the network model.

**Fig 7.**
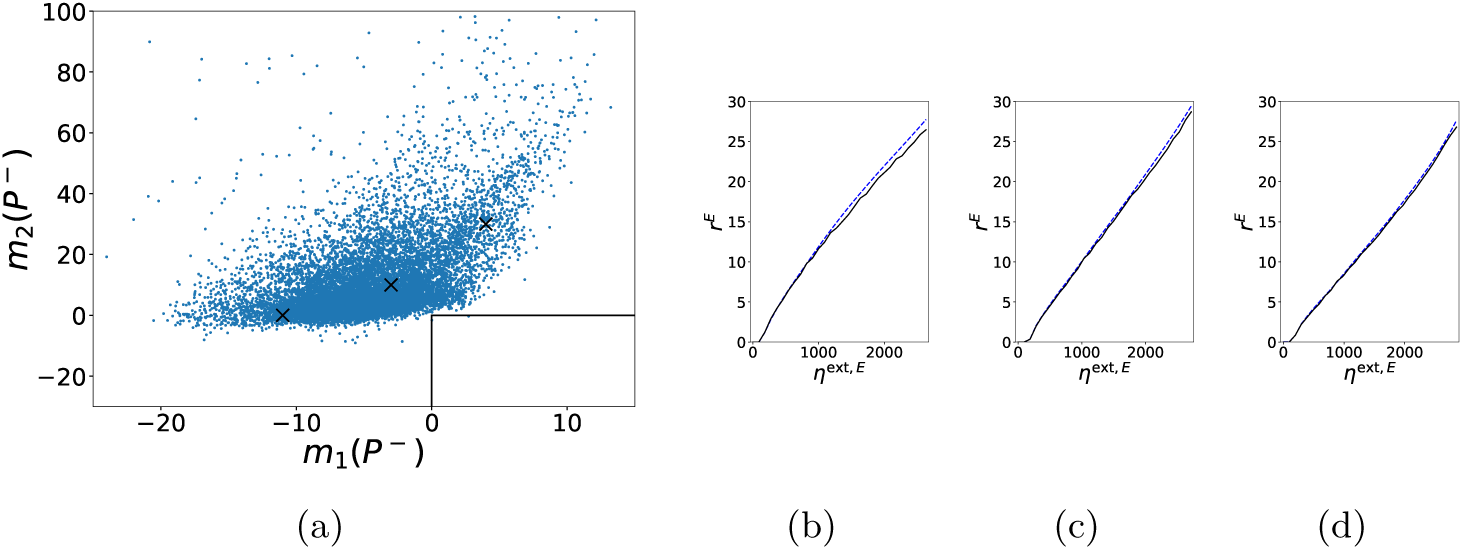
Model capability in generating sigmoidal gain curve. (a) *m*_1_ vs. *m*_2_ (see main text for notation) for physiologically plausible *P* ^−^’s. (b), (c) and (d) are example gain curves corresponding to the three typical data points [−11, 0], [−3, 10] and [4, 30] marked in (a), respectively. Blue dashed lines indicate surrogate gain curves 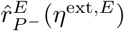 whereas black lines indicate the simulated gain curves 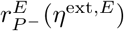.

We conclude that the integrate-and-fire model described in **Materials and** methods (I&F neuronal model) without further enhancement is incapable of producing gain curves that are sigmoidal in shape and that deviate substantially from a straight line. This is a limitation of the model. The present study should serve to inform the modeling community that to produce a sigmoidal gain curve with more pronounced curvature (as has been observed experimentally), some other mechanisms must be incorporated. In the V1 network model in [38], for example, mechanisms such as synaptic depression of I-neurons and potassium currents that prevent E-cells from firing repeatedly in rapid succession were implicated in contrast response properties.

## Discussion

A broader aim of this work is to promote the use of machine-learning approaches in biological modeling. We propose that these more systematic methods can be useful not as replacement of but as supplement to conventional modeling techniques [12]. To demonstrate the efficacy of this approach, we considered a neuronal network built to resemble local circuits in the cerebral cortex, and illustrated how via the use of a surrogate DNN combined with data analysis (such as correlation analysis and logistic regression), rich statistical structures can be extracted from limited data generated by simulation.

A specific approach that we are proposing here is the following: While biological processes are typically extremely complex, if one is able to build a model of the system modulo a finite – possibly very large – number of unknown parameters and identify a finite number of key quantities that best describe what goes on, then the modeling problem can be framed in terms of discovering the mapping from

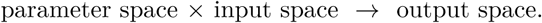

Such input-output relations are especially well suited to data-driven inferences using neural nets. The statistical analysis of DNN surrogates in general suggests rather than prove any specific behavior of the target mapping, due to the presence of uncertainties intrinsic to any data-driven approach. Nevertheless, compared to heuristic arguments and *ad hoc* numerical explorations of parameter space, these results are quantitative in nature and provide strong supporting evidence for the conclusions they suggest.

### On surrogate-based modeling and DNN

After a surrogate learns from data, it allows highly efficient manipulation including evaluation, differentiation, optimization (e.g. parameter tuning) and statistical analysis. Among a rich class of conventional surrogate models, many of which may serve our purpose equally well, DNN is convenient to use for a number of reasons: there are rich and sophisticated open source libraries (e.g., Tensorflow, Keras, Pytorch); DNN is faithful to data, with low training error; it is robust, generalizes well, and often does not require extra regularization; finally it is flexible, with universal approximation capability and rich architecture.

In engineering, the use of surrogate-based modeling to assist in the analysis and exploration of complex experiments and designs is well established. On the other hand, in spite of its huge success in tasks related to image, audio, and video recognition and processing, DNN up until now has largely remained a black box. It is only in recent years that researchers from different scientific disciplines have begun to exploit its many potentials. We believe that DNN surrogates can potentially be of great use in biological modeling, and it is with more complex models in mind that we have embarked in this direction. This paper is a first step to demonstrate, using a network model of a local cortical circuit, the type of statistical analysis made possible by such an approach.

### Applications to neuroscience

Many questions remain. For local circuits, the *P* in our *P* → *O* mapping can include, e.g., connectivity and system size, *O* can be currents, and an important problem inspired by the balanced-state ideas in [39–42] may be to quantify the balancing of currents under different network conditions. Nor must the target *O* be limited to firing rates and currents. It can include other quantitative measures of firing patterns, such as correlations and degrees of synchrony. A problem of interest is to relate gamma rhythms as characterized by their power spectral densities to network parameters [43, 44], as gamma rhythms are known to be altered by disease, drugs and other physiological states [45–47] These are all potential applications of the methodology proposed.

Populations of homogeneously connected and homogeneously driven neurons are ideal starting points for theoretical studies. A natural next step is to consider multi-component networks, beginning with source-target populations and progressing to more complicated network motifs with feedback loops. Neuronal networks in the real cortex are in fact not abstract graphs; they have spatial structures (see e.g. [48]). An ultimate use of DNN-assisted surrogates may be to reveal the mapping

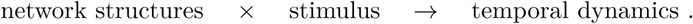

### Outlook on the use of surrogates in biological (neural) modeling

High degrees of complexity and a low ratio of knowns to unknowns is characteristic of biological modeling. A case in point is the modeling of neuronal circuits. Network models that incorporate neuroanatomy and physiology are necessarily very complicated because of the large numbers of neurons (on the order of 10^11^ in the human cerebral cortex), the many neuron types, their detailed and varied modes of interactions, not to mention the complex wiring, with intra/inter-laminar connections, and inter-areal connections with multiple feedforward and feedback loops.

This level of complexity implies (i) any realistic model will contain a large number of unknown parameters; (ii) *a priori* constraints for many of these parameters are hard to obtain, and (iii) simulation time is long, limiting the number of training sets possible. The issues above exacerbate one another. For example, when parameter space has dimension *d* » 1, a search domain that is *k* times larger in each dimension will result in a volume that is *k*^*d*^ times larger; and if the actual physiological domain is small relative to the search domain, then with high probability, a reasonable-sized sample will not contain a single point in the actual physiological domain.

In Fig. 8 we used a parameter domain 𝒫_L_ with *k* ≈ 5 compared to 𝒫, the domain used in Results. Using a training set of 40000 points sampled randomly from 𝒫_L_, it was very likely that none was 𝒫. This figure shows, however, our well-trained DNN still achieved a good accuracy of ∼1Hz. Compared to Fig. 1(b), a larger training set was needed, and the accuracy was lower, but it performed satisfactorily nevertheless.

**Fig 8.**
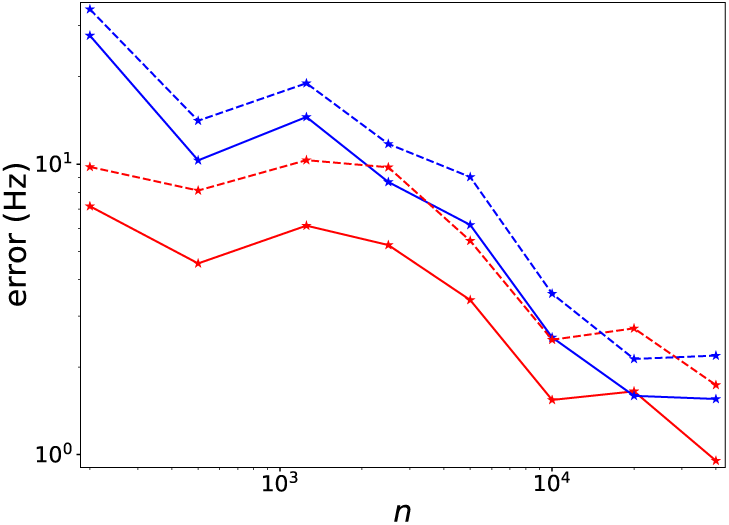
Performance of DNN trained on a large domain. Performance of DNN trained by training datasets of size *N* sampled from 𝒫_L_ tested on a test dataset of size 10000 sampled from 𝒪 is presented. MAE (dashed line) and RMSE (solid line) for *r*^*E*^ (red) and *r*^*I*^ (blue) are exhibited.

In **Results** (**Viability of parameters and DNN performance**) and again in Fig. 8, the reason why small training sets sufficed was the simplicity of the mapping from input to output, a fact we confirmed in subsequent sections. Obviously one cannot conclude from this one study that such mappings always have simple structures, but modeling experience of the authors suggests that even in large-scale biologically realistic network models (e.g. [38]) neuronal responses tend to depend fairly smoothly on parameters. This means that locally in parameter space, the dependence of target mappings on parameters is relatively simple, not unlike those revealed in our derivative analysis.

These observations offer hope to the feasibility of surrogate-based approaches for more complex neuronal circuit models. They also point to the need for good *a priori* bounds on physiological ranges to help simplify the structure of input-output maps, and this is where biology enters. The judicial use of biological facts and experimental data to partially constrain parameters in advance will increase the chances of success for machine-learning approaches.

## Materials and methods

We first describe the neuronal model that was used for illustration throughout the paper. Then, we define the deep neural network that was used as surrogate for this model. At last, we briefly introduce correlation analysis and logistic regression.

### I&F neuronal model

In this work, we consider a homogeneously connected network of integrate-and-fire (I&F) neurons that can be thought of as a generic model of a local neuronal population. The network has *N*_*E*_ = 225 excitatory neurons (E-neurons) and *N*_*I*_ = 75 inhibitory neurons (I-neurons) with a ratio of *N*_*E*_*/N*_*I*_ = 3. Each E-neuron is postsynaptic to another E-neuron with probability 10% and to an I-neuron with probability 50%. Each I-neuron is postsynaptic to any other neuron with probability 50%. These connection probabilities are consistent with those in the visual cortex; see [7] for supporting references. A single realization of the random graph with these connectivities was fixed and used throughout in our numerical experiments.

The dynamics of each neuron in the network is modeled by the I&F equation

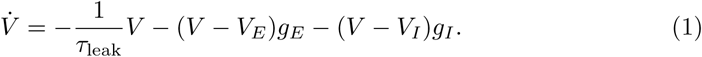

Here time is in milliseconds (ms) and *V* is the membrane potential normalized in a dimensionless unit with a reset value *V*_*R*_ = 0 and a spiking threshold *V*_*T*_ = 1, so that when *V* reaches *V*_*T*_, the neuron fires a spike; then *V* is reset to *V*_*R*_ and will remain there for an absolute refractory period of 2.5ms. In these normalized units, *V*_*E*_ = 14*/*3 and *V*_*I*_ = −2*/*3 are excitatory and inhibitory reversal potentials, and *τ*_leak_ = 20ms is the leak rate [49]. For any neuron *n* of type *Q* ∈ {*E, I*}, *g*_*E*_, *g*_*I*_ *≥* 0 are its excitatory and inhibitory conductances governed by

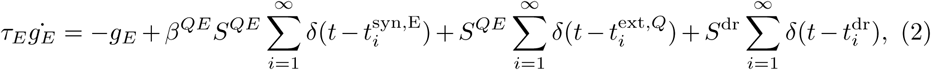

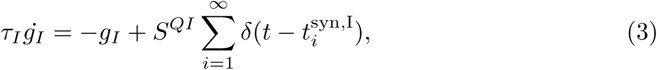

where *τ*_*E*_ = 2ms and *τ*_*I*_ = 3ms are decay rates for excitatory and inhibitory conductances respectively. Synaptic inputs from other neurons within the network are described in the second terms on the right sides of Eqns (2) and (3): 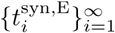 and 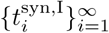 are the spike times of all the E- and I-neurons presynaptic to neuron *n*, and *δ*(·) is the dirac delta function indicating an instantanous jump of conductance *g*_*E*_ or *g*_*I*_ upon the arrival of an E or I-spike, with strength/amplitude equal to *β*^*QE*^*S*^*QE*^*/τ*_*E*_ and *S*^*QI*^*/τ*_*I*_ respectively. The quantity 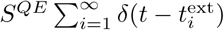 models the independent excitatory drive to neuron *n* from another region of the brain with Poisson kicks at rate *η*^ext,*Q*^ arriving at times 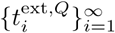. In addition, neuron *n* receives an independent Possion drive with strength *S*^dr^ = 0.005, rate *η*^amb^ and arrival times 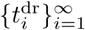; this term is intended to represent “ambient” modulatory influences from other parts of the brain or body. Note that we do not model synapses individually, and to simulate the effect of synaptic failure between E-neurons, at each spike a random number *β*^*EE*^ is picked from the uniform distribution on [0.8, 1]; we have set *β*^*IE*^ = 1, i.e., no synaptic failure for the synapses from E- to I-neurons is assumed.

The undetermined parameters of this model are the synaptic coupling weights among model neurons, *S*^*EE*^, *S*^*EI*^, *S*^*IE*^ and *S*^*II*^, and inputs parameters to the population *η*^ext,*E*^, *η*^ext,*I*^ and *η*^amb^.

Synaptic weights of real cortical neurons are not known, but physiologically plausible ranges can be estimated from a combination of indirect measurements (such as *in vitro* experiments and the firing rates of neurons) together with some analysis (see [7], Methods). In this paper, we will assume the physiologically plausible ranges to be

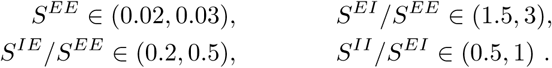

We have chosen to normalize the other quantities by *S*^*EE*^ because it has been observed from parameter tuning (in e.g. [7]) that the 4 synaptic weights 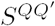 can be adjusted up and down together without having a strong effect on the system; this point will be justified later on in our analysis. Note that *S*^*II*^ is normalized by *S*^*EI*^ with a ratio less than 1 to account for electrical coupling among I-neurons, which effectively weakens the self-inhibition of the I-population.

With regard to the input parameters, in this paper we will assume the plausible ranges are

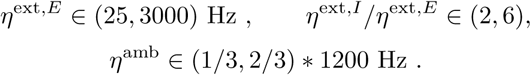

The range for *η*^ext,*E*^ is large as it is intended to include input strengths that range from spontaneous to strong drive, and we have coupled the drive to E and to I-neurons because most synaptic input will affect both. The quantity *η*_0_ = 1200 Hz is the threshold for causing a neuron to spike in the absence of other inputs, and *η*^amb^ in real cortex is known to be below this threshold.

From here on, we will refer to the parameters above as *P* = [*P*_S_, *P*_I_], where

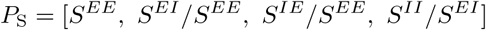

are network synaptic parameters and

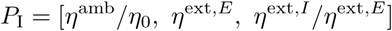

are input parameters, and we will say *P* = [*P*_S_, *P*_I_] is in our physiological domain 𝒫, if all 7 parameters fall within the ranges above.

Given *P*, we let *r*^*E*^ and *r*^*I*^ denote the mean firing rates of the E- and I-populations at steady state, and our model output is taken to be

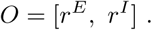

Model firing rates are computed through numerical simulation. In our simulations, each trial runs for 3s, the last 2s of which are used to compute the system’s (empirical) firing rates. We assume, based on physiological experiments, that in an active state of the cortex,

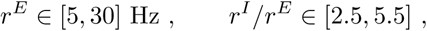

and we will say *O* is in our physiological domain 𝒪 if both *r*^*E*^ and *r*^*I*^ fall in the ranges above.

We reiterate that 𝒫 consists of *a priori* biological constraints either deduced from indirect experimental measurements or learned from previous modeling results. It is necessary to partially constrain parameter space, and these are effectively educated guesses. The domain 𝒪, on the other hand, consists of firing rates that correspond roughly to what is observed in the laboratory under a variety of circumstances. There is no guarantee whatsoever that *P* ∈ 𝒫 will produce *O* ∈ 𝒪.

### DNN surrogate

First we review the general setup for a DNN. For the regression problem of fitting a training dataset 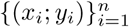, where *x*_*i*_ ∈ ℝ^*d*^ and 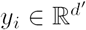 for each *i*, a fully connected DNN of *H* layers, *H ≥* 2, is defined as follows. Let 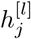 be the output of the *j*th node of the *l*th layer of the DNN. Then

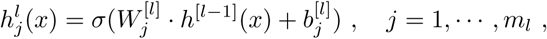

where *x* ∈ ℝ^*d*^, *m*_*l*_ is the number of neurons in layer 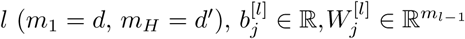, and 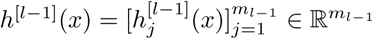. For the *j*th neuron of the output layer of the DNN,

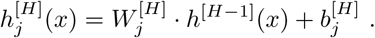

The DNN is abbreviated as *h*(*x*; *θ*) = *h*^[*H*]^(*x*), where

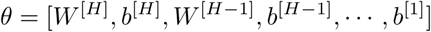

is the set of parameters of the DNN. In this work, the activation *σ* is fixed to the sigmoid function, i.e., 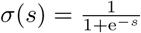. The loss function is fixed to the mean-square error (MSE)

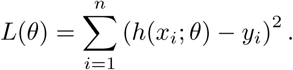

During training, the parameters of the DNN in each epoch *t* can be updated using gradient descent as

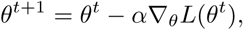

where *α* is the learning rate. To speed up the training process, we use a popular accelerated gradient-based optimizer of Adam in our experiments [24].

Here is how the DNN will be used in this work: We train a sigmoid-DNN *h*(*x*; *θ*) of hidden layer sizes 800 − 200 − 200 on training dataset 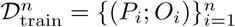 obtained from *n* trials of simulations (for various values of *n*), where each *P*_*i*_ is randomly drawn from a uniform distribution in its physiological domain 𝒫. The accuracy of the DNN *h*(·; *θ*_*n*_), where *θ*_*n*_ is the weight of DNN well-trained on 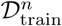, is evaluated on a testing dataset 𝒟_test_ consisting of 10000 (*P, O*)-pairs where *P* was drawn independently from and *O* was computed from simulations. Mean-absolute error (MAE) defined as 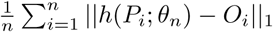 and root-mean-square error (RMSE) defined as 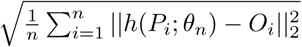 are used for accuracy quantification. A DNN trained on 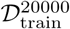, donoted by 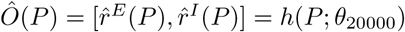, serves as a surrogate of the neuronal circuit for all later analysis.

We remark on the following known properties of the DNN that make it a powerful tool: (i) DNN is a universal approximator. It has been proved that a sufficiently wide neural network of at least one hidden layer can approximate any continuous function to any desired accuracy [50–52]. (ii) Empiricial and theoretical studies indicate that the DNN approach is free from the curse of dimensionality, i.e., error decay can be bounded by a scaling 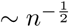 independent of the input dimension [18, 19]. (iii) It has been observed in practice that DNNs in general do not overfit even in an overparameterized setting without explicit regularization [53]. Non-overfitting combined with the universal approximation property makes DNN a highly robust and flexible approach for capturing general nonlinear mappings. (iv) It has been shown by the discovery of the Frequency Principle that DNNs are especially effective in learning low frequency functions from training data [20, 21, 54, 55]. Therefore, very good accuracy can be achieved if the target mapping is dominated by low frequencies.

Finally, as noted in the Introduction, other machine learning approaches like support vector regression (kernel method) and gaussian process regression (kriging) may also serve our purposes of surrogate modeling. However, we anticipate that, for more complex biological networks, the flexibility of DNN surrogates may be a great advantage in application.

### Correlation analysis and logistic regression

Correlation between two variables *x*_*i*_, *x*_*j*_ *∈ {*−1, 1} is defined by

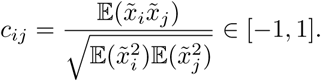

where 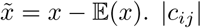 is also a good indicator of how accurate *x*_*i*_ and *x*_*j*_ can predict one another.

Logistic regression solves a classification problem as follows. Model 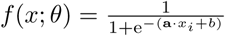 with *θ* = [**a**, *b*] is fitted to data 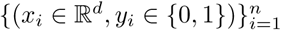 by maximizing the log-likelihood function, i.e.,

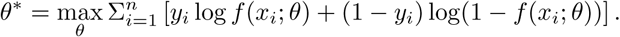

Then, for any *x*, if *f* (*x*; *θ*^*\**^) *>* 0.5, the output is predicted as 1, otherwise as 0. A high prediction accuracy indicates that input domains correspond to different outputs are linearly separable, whereas low prediction accuracy (≲50%) indicates a complex structure not linearly separable. In Results, to use logistic regression for the prediction of sign of derivatives, we map positive sign to 1, negative sign to 0 and solve the optimization problem above.

## Acknowledgments

Yaoyu Zhang was supported by NSF Grant No.DMS-1638352 and the Ky Fan and Yu-Fen Fan Membership Fund. Lai-Sang Young was supported in part by NSF Grants 1734854 and 1901009. The authors would like to thank David Hansel, Aaditya Rangan and Robert Sharpley for valuable comments and suggestions.

